# A generalizable multivariate brain pattern for interpersonal guilt

**DOI:** 10.1101/835520

**Authors:** Hongbo Yu, Leonie Koban, Luke J. Chang, Ullrich Wagner, Anjali Krishnan, Patrik Vuilleumier, Xiaolin Zhou, Tor D. Wager

## Abstract

Feeling guilty when we have wronged another is a crucial aspect of prosociality, but its neurobiological bases are elusive. Although multivariate patterns of brain activity show promise for developing brain measures linked to specific emotions, it is less clear whether brain activity can be trained to detect more complex social emotional states such as guilt. Here, we identified a distributed Guilt-Related Brain Signature (GRBS) across two independent neuroimaging datasets that used interpersonal interactions to evoke guilt. This signature discriminated conditions associated with interpersonal guilt from closely matched control conditions in a cross-validated training sample (*N* = 24; Chinese population) and in an independent test sample (*N* = 19; Swiss population). However, it did not respond to observed or experienced pain, or recalled guilt. Moreover, the GRBS only exhibited weak spatial similarity with other brain signatures of social affective processes, further indicating the specificity of the brain state it represents. These findings provide a step towards developing biological markers of social emotions, which could serve as important tools to investigate guilt-related brain processes in both healthy and clinical populations.

## Introduction

Guilt is an experience that arises when we violate norms or values that we consider important—for example, when we have wronged someone whose wellbeing we care about. Guilt is considered a quintessential moral emotion, as it plays a crucial role in motivating adherence to social norms and promoting conciliation after interpersonal conflict (Baumeister et al. 1994; Hoffman 2001; Tangney et al. 2007; Tooby and Cosmides 2008; Sznycer 2018). It is also a core feature of several important clinical conditions. On the one hand, a lack of guilt is a central feature of psychopathy, and is associated with antisocial behavior (Viding et al. 2009; Blair 2013). On the other hand, depression, suicidal ideation, and other internalizing disorders are associated with excessive guilt (Tilghman-Osborne et al. 2012; Ratcliffe 2014). Understanding how the brain represents this complex moral emotion can then inform the development of translational applications to clinical settings (Huys et al. 2016).

Emotion theorists have proposed that guilt arises from a particular type of cognitive appraisal that includes several elements: (1) the recognition that one’s actions or inaction is causing suffering, (2) affiliation or identification with the suffering other (e.g., a friend or other ingroup member), and (3) attribution of blame or responsibility to oneself (i.e., one could have acted differently) (Frijda 1993; Baumeister et al. 1994; Chang and Smith 2015). Although appraisal theory suggests that guilt arises from a unique set of appraisals with, potentially, a unique ‘constellation’ of brain ingredients (Moors et al. 2013), such patterns need not necessarily be mapped to brain features in a consistent way across instances of guilt and across individuals (Barrett and Satpute 2013). Thus, it remains unclear as to whether there is a reliable ‘signature’ associated with this particular configuration of thoughts and beliefs, and whether such processes have consistent brain responses across individuals and tasks. Being able to identify such stable signature of guilt could inform us about how similar or dissimilar the neural processes related to guilt are to those underlying other affective states (e.g., sadness, regret, etc.). In addition, identifying stable guilt-related brain signature would be an important step towards understanding the function and dysfunction of the underlying brain circuitry in healthy and clinical populations (Woo et al. 2017).

Two recent neuroimaging studies have manipulated guilt in interpersonal interactions by manipulating the two key guilt-related appraisals — (1) perception of others’ suffering, and (2) the knowledge that one’s actions caused that suffering (Koban et al. 2013; Yu et al. 2014). These studies have shown that both features are determinants of self-reported guilt (but not other emotions; Koban et al. 2013) and consistently found that they are associated with increased activation of anterior/middle cingulate cortex (ACC/aMCC) and bilateral anterior insula (AI). However, the univariate analyses adopted in these studies are not sufficient to provide a brain signature of guilt. First, the univariate approach seeks each single voxel that shows significant difference in activation strength across different psychological states. The differences in psychological states, however, may not be encoded in the activation strength of any single voxel; rather, it may be encoded by distinct patterns of activation across a large number of voxels (or the whole brain). Second, the univariate analysis is designed to test for nonzero correlations between psychological states and brain measures, but not to estimate predictive accuracy (effect size) of the identified brain correlates. For example, although both Yu et al. (2014) and Koban et al. (2013) reported the activation of the aMCC and anterior insula in high relative to medium or low guilt conditions, the activation in each study cannot be used to predict experimental conditions in the other study, rendering it difficult to conclude whether the activations in the two studies are reliably similar.

Here, we address these open questions and develop a neurophysiological signature of guilt-related cognitive appraisals. To be clear, we do not treat the signature as *the* necessary and sufficient neurophysiological conditions for guilt, namely, capturing all and only neurophysiological states associated with guilt. However, it is still useful as a provisional marker that confers information value, as well as a defined brain measure, for provisional inference, comparisons, and further testing and validation on the brain bases of social emotions (Kragel et al. 2018). Such a neural signature should satisfy three criteria: sensitivity, specificity and generalizability (Woo and Wager 2015; Krishnan et al. 2016; Woo et al. 2017). Specifically, it should: 1) detect the presence of the *cognitive antecedents* of guilt (i.e., sensitivity); 2) not respond to negative experiences elicited by other affective stimuli, such as physical pain and general negative affect (i.e., specificity); and 3) generalize across studies and samples where the cognitive antecedents (i.e., not necessarily the subjective feelings) on which the signature is trained are present (generalizability).

To achieve this aim, we used a predictive modeling approach to identify a whole-brain pattern that is sensitive and specific to the core antecedent of guilt—being causally involved in harming others during interpersonal interaction (Koban and Pourtois 2014; Cui et al. 2015). We adopt an analytic approach (Kragel et al. 2018) that has been successfully applied to investigating the neural representation of various affective processes, including physical pain (Wager et al. 2013), vicarious pain (Krishnan et al. 2016), social rejection (Woo et al. 2014), unpleasant pictures (Chang et al. 2015), basic emotions (Lindquist and Barrett 2012; Kragel and LaBar 2015; Wager et al. 2015; Kragel et al. 2016; Saarimäki et al. 2018), empathy (Ashar et al. 2017) and related social emotions (Saarimäki et al. 2018). We trained a support vector machine classifier to discriminate brain states elicited by social contexts that differ only in one’s causal role in the other’s suffering (Study 1; Yu et al. 2014). We then tested the model’s generalizability by applying the obtained multivariate guilt pattern to a second neuroimaging dataset, which employed a similar interpersonal action-monitoring paradigm in a different population and using a different MRI scanner (Study 2; Koban et al. 2013). Further convergent and discriminative validity of the pattern was assessed by examining its performance in predicting subjective guilt ratings and compensation behavior—participants’ willingness to voluntarily accept painful shocks in order to reduce further shocks administered to a person they believe they harmed—and testing specificity against several other negative affective states (e.g., physical pain, vicarious pain, emotion recall). Altogether, datasets from 4 independent studies (*N* = 86 healthy participants) were used for training and testing the signature.

## Materials and methods

### Participants

For Study 1, twenty-four undergraduate and graduate students (mean age 22.0 years; 11 female) were recruited at the Southwest University, Chongqing, China (Yu et al. 2014). Nineteen adults (mean age 24.3 years; 9 female) participated in Study 2, conducted in Geneva, Switzerland (Koban et al. 2013). All participants in the final sample (total of *N* = 43) had normal or corrected-to-normal vision and none reported any history of psychiatric or neurological disorders. All participants provided informed consent before scanning and were paid for their participation.

### Procedure

Both Study 1 and Study 2 adopted an interactive paradigm where a participant in the scanner and a participant outside the scanner performed a dot-estimation task that involved estimating the number of dots briefly presented on a screen (for similar interactive paradigms, see Kédia et al. 2008; Cui et al. 2015; Lepron et al. 2015). Mistakes in the dot-estimation task would result in the out-of-scanner participant (hereafter, “partner”) receiving mildly painful stimuli. Essentially, both studies manipulated participants’ responsibility for the harm to the partner. In Study 1 (Fig. 1A), participants underwent two fMRI scanning blocks. In the first block (i.e., Pain block), participants were told that the partner (confederate) would receive mild electric shocks if either the partner, the participant, or both made a mistake in a dot-estimation task. This allowed us to manipulate increasing levels of guilt, with some guilt expected whenever a mistake occurred (“Pain: Partner_Responsible”, “Pain: Both_Responsible”), and the most guilt when the participant, but not the partner, responded incorrectly (“Pain: Self_Responsible”). On the trials where the partner would receive electric shocks, the participants were given the option to intervene and bear a proportion of pain for the partner. In the second block (i.e., NoPain block), the participants were told that they would interact with the same partners in an almost identical task, with the exception that no pain stimulation was delivered to either side. The NoPain block was included as a guilt-free control for psychological processing of correct/incorrect feedback and the process of making social comparisons (i.e., comparing one’s own performance with the partner’s performance).

**Figure 1.**
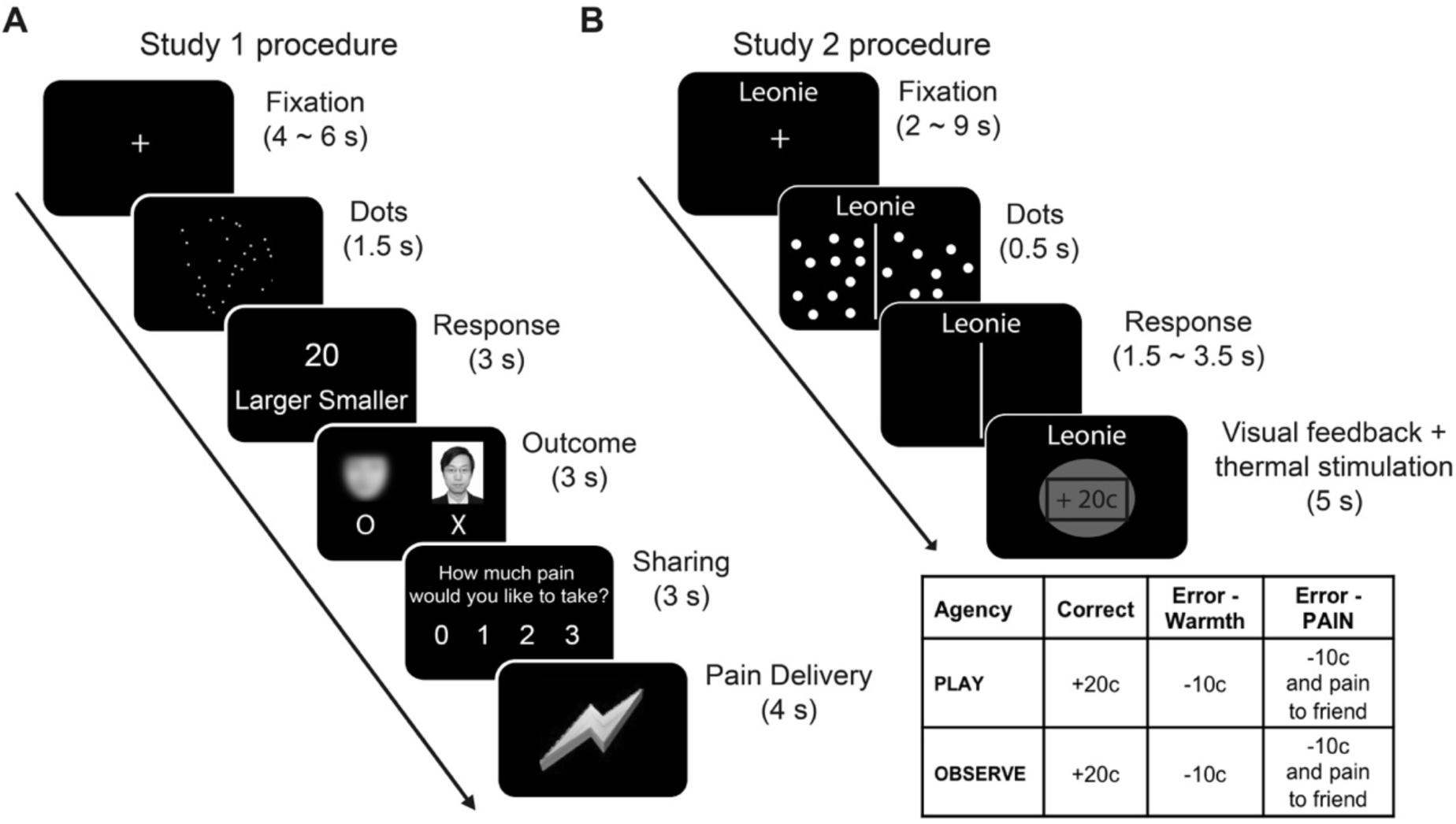
Procedure for Study 1 and Study 2. (A) In Study 1, the participant in the scanner was randomly paired with an anonymous partner on each trial. The task for the participant and the partner was to quickly estimate the number of dots presented briefly on the screen. The feedback of their estimations was presented under the photo of the participant and under a blurred picture of face representing the partner. If at least one of them estimated wrong, the partner need receive a number of mildly painful electric shocks. The participant then indicated the level of pain he/she would be willing to take for the partner Finally, a pain stimulation of the participant’s choice was delivered to him/her (see ref (Yu et al. 2014) for details). (B) In Study 2, two participants took turns in either performing or observing the other’s performance in a dot estimation task. The dot estimation task required both participants to indicate which side of the screen contained greater number of dots. The participant outside the scanning room would receive either painful or nonpainful (i.e., warm) thermal stimulation after each trial, depending on the performance of the current player. The full 2 × 3 factorial design resulting from the different feedback type in the two task conditions (playing or observing) is displayed in the table (see ref (Koban et al. 2013) for details).

In Study 2 (Fig. 1B), participants played a similar dot estimation task. In alternating blocks, the participant in the scanner (i.e., Play block) took turns with an actual friend who was situated outside the scanner (i.e., Observe block) to perform the dot estimation task. Both friends would win points for correct responses and lose points (later converted into bonus money) for erroneous (incorrect) responses made by either player (Play or Observe condition). Crucially, the participant outside the scanner would receive additional painful heat stimulation on a randomly selected half of the error trials and non-painful warmth stimulation on the other half of the error trials, and were informed when the partner was receiving pain. This resulted in a 3-by-2 factorial design with three levels of Feedback (Correct, Error_Warmth, and Error_Pain) and two levels of Agency (Play vs. Observe). It was expected that the condition in which the participant inside the scanner caused pain to a friend by making an error (i.e., Play: Error_Pain) would lead to the highest levels of guilt.

### Post-scan manipulation check (emotion ratings)

After scanning, an emotion manipulation check was employed in both studies. In Study 1, participants rated their feelings of guilt, fear, anger, and distress, for each of the 3 experimental conditions, in which an incorrect response occurred. In Study 2, participants rated, their feelings of guilt, fear, anger, shame, and sadness, for each of the 6 experimental conditions.

### Neuroimaging Data Acquisition

For Study 1, images were acquired using a 3.0-Tesla whole-body scanner (Trio TIM, Siemens, Germany). T2*-weighted functional images were acquired in 36 axial slices parallel to the AC–PC line with no interslice gap, affording full-brain coverage. Images were acquired using an EPI pulse sequence, with a TR of 2200 ms, a TE of 30 ms, a flip angle of 90°, an FOV of 220 mm × 220 mm and 3.4 mm × 3.4 mm × 3.5 mm voxels. A high-10 resolution, whole-brain structural scan (1 × 1 × 1 mm^3^ isotropic voxel) was acquired after functional imaging. For Study 2, images were acquired using another 3.0-Tesla whole-body scanner (Trio TIM, Siemens, Germany). T2*-weighted EPI sequence (2D-EP, repetition time = 2100 msec, echo time = 30 msec, flip angle = 80°, 3.2 × 3.2 × 3.2 mm^3^ voxel size) for acquisition of functional images of the whole brain (36 slices). The structural image of each participant was recorded with a T1-weighted MPRAGE sequence (repetition time = 1900 msec, inversion time = 900 msec, echo time = 2.27 msec, 1 × 1 × 1 mm^3^ voxel size).

### Neuroimaging data analyses

#### Preprocessing and univariate general linear model (GLM) analyses

Details of preprocessing are described elsewhere (Yu et al. 2014 for Study 1; Koban et al. 2013 for Study 2). In brief, univariate general linear model (GLM) analyses were conducted in SPM8. For both studies, the critical regressors were those corresponding to the feedback of the visual task. For Study 1, trials from the Pain block and the NoPain block were modeled in separate GLMs. Each model contained as critical regressors the following conditions, all modeled with HRF starting at the onset of the feedback of the dot-estimation task and covering the entire feedback phase (duration = 3 secs): the condition in which the participant alone made a wrong response (“Pain: Self Responsible”), the condition in which both players made a wrong response (“Pain: Both_Responsible”), the condition in which the partner alone made a wrong response (“Pain: Partner_Responsible”), and the condition in which both players made a correct response (“Pain: Both_Correct”). Also included were regressors of no interest: cue for new trial, random dot presentation, estimation responses, compensation responses and pain delivery (the last two were only included for the Pain block). For Study 2, the relevant regressors corresponded to the feedback in the six experimental conditions: Error_Pain, Error_Warmth, and Correct in both the Play block and the Observe block. The contrast images corresponding to the main effects of these regressors versus baseline were extracted and used for training and test in the multivariate pattern analysis.

#### Guilt pattern classification

We trained a linear support vector machine (SVM; slack parameter C = 1 was chosen *a priori*) to discriminate “Pain: Self_Responsible” (high guilt; coded as 1 in the classification) versus “Pain: Both_Responsible” (medium guilt; coded as −1 in the classification) conditions in Study 1 with a leave-one-subject-out cross-validation procedure (Friedman et al. 2001; Wager et al. 2013; Woo et al. 2014). The rationale of training the classifier to discriminate these two conditions is to avoid as much as possible the classifier capturing processes that are not essential for detecting guilt. For example, a classifier trained to discriminate the “Pain: Self_Responsible” and the “Pain: Both_Responsible” conditions would not only capture the responsibility of the participant in causing pain, but would also capture differences in the outcome feedback of participant’s performance (i.e., correct vs. incorrect). In the statistical learning literature (Friedman et al. 2001), there are many types of classification algorithms, but they generally perform very similarly on problems such as the one we pursued here. Support vector machine algorithms such as the one we used in this study are the most widely used algorithm for two-choice classification, and are robust and reasonably stable in the presence of noisy features. Exploring different algorithms could be interesting, but may lead to an open-ended, largely methodological pursuit that is not expected to impact performance in a reproducible or systematic way in the present datasets. In addition, we wanted to avoid the trap of fitting multiple algorithms and picking the best one, thus overfitting the dataset. Therefore, we chose a widely used algorithm (whole-mask SVM) whose effectiveness has been well established in previous studies.

The images used in this analysis were the whole-brain activation maps masked by an *a priori* meta-analytic map associated with the term ‘Emotion’ from Neurosynth (uniformity test map, thresholded at *p*_FDR_ < 0.01, accessed as of September 7^th^ 2014, see Fig. S1 for details; Yarkoni et al. 2011). This mask was chosen to select voxels that are presumably important for emotional processing in the brain. We acknowledge that emotions are likely emergent processes from interactions between many brain regions (Scherer 2009; Lindquist et al. 2012; Pessoa 2017), potentially including those outside typical ‘emotion’ brain regions captured by this Neurosynth mask. However, there is a trade-off between considering all possible features (i.e., voxels) and generalizability of the classifier across participants and studies, because a classifier may pick up on noisy dimensions that do not generalize well to new datasets. Thus, the (methodological) rationale of applying the Emotion mask prior to classifier training was feature selection and dimension reduction, with the aim of decreasing overfitting and of increasing generalizability. We assume that although Study 1 and Study 2 induced guilt with slightly different interactive tasks, the core underlying emotional processes should overlap. Training and testing the classifier within the emotion mask could reduce the possibility of overfitting and therefore increase the generalizability of the classifier to the test dataset. Future research with larger sample sizes could investigate the role of other areas in the brain and use nested cross-validation for optimizing feature selection and the trade-off with generalizability.

The procedure trains the classifier on N-1 participants and generates a weight map that best classifies the sample, and tests the classification on the left-out (N^th^) participant. This process is repeated until all participants have served as the test sample for the classification algorithm exactly once to obtain their respective cross-validated signature response values. The classifier obtained thus represents a hyperplane in the feature space that best separate the observations (i.e., individual brain activation maps) in the “Pain: Self_Responsible” condition and the “Pain: Both_Responsible” condition.

#### Guilt pattern expression

The contrast images from the first-level analysis for each participant were used to obtain pattern expression values for the guilt pattern. To obtain single pattern expression values for each condition and each participant, we computed the dot product of the cross-validated weight map of the guilt pattern and the individual contrast images. This value reflects the distance between a given activation map and the classifier represented by a hyperplane represented in the feature space. These pattern expression values were then tested for differences between experimental conditions. We calculated the forced-choice classification accuracy for how well the two conditions in questions were correctly classified based on their pattern expression values. A sensitive and generalizable pattern for interpersonal guilt should be not only able to discriminate the “Pain: Self_Responsible” versus “Pain: Both_Responsible”, on which the classifier was trained, but also to separate the “Pain: Self_Responsible” and other less guilty conditions in Study 1 (i.e., Pain: Partner_Responsible and Pain: Both_Correct), as well as different guilt states in Study 2. Additionally, the pattern expression values for the conditions in the Pain block of Study 1 were regressed against the willingness to accept the partner’s pain in respective conditions to assess their ability in predicting guilt-induced compensation behavior. In the regression model, condition was included as a dummy variable to covariate out the variation of compensation as a function of conditions.

For specificity, the predictive power of the interpersonal guilt pattern should not generalize to other types of negative affect. To test the specificity of the guilt pattern, we obtained individual activation maps for unpleasant experiences other than interpersonal guilt, including physical pain and vicarious pain (Study 3, *N* = 28; Krishnan et al. 2016), and emotion-recall (Study 4, *N* = 15; Wagner et al. 2011). Study 3 dataset contained three sets of maps corresponding to three levels of thermal pain (high, medium, low) applied on the volar surface of the left forearm and three sets of maps corresponding to viewing three levels of unpleasant images (high, medium, low). The emotion-recall dataset contained three sets of maps corresponding to participants’ recall of personal memories of past experiences of guilt, sadness, and shame.

#### Comparison with other brain signatures

To investigate the spatial similarity of the guilt signature with other patterns (masked by the same Emotion meta-analytic map as the GRBS), we calculated the spatial similarity (Pearson correlation coefficient) between the GRBS and signatures for physical pain (NPS, Wager et al. 2013), picture-induced negative affect (PINES, Chang, et al, 2015), social rejection (Woo et al. 2014), vicarious pain (VPS, Krishnan, 2016), empathic distress and empathic care (Ashar et al. 2017) and skin conductance and heart rate (Eisenbarth et al. 2016)

Further, we investigated the local pattern similarity of the GRBD and the PINES within the meta-analytic Emotion-mask and within three canonical emotion-related brain regions—ACC, insula, and amygdala. We used enhanced scatter plots (Koban et al. 2019) to visualize the amount of shared positive, shared negative, and unique positive and negative voxel weights for two signatures (*z*-scored to make them comparable) in those areas. As described in detail before (Koban et al. 2019), each voxel’s weights for the two signatures were plotted on the x- and y-axis respectively, and this scatter plot was then divided into eight sectors (octants), reflecting different directions of shared and unique weights for each pattern. Voxels in Octant 1 had positive weights for the GRBS, but near-zero weights for the PINES, voxels in Octant 2 had positive weights for both patterns (reflecting shared variance), voxels in Octant 3 had positive weights for the PINES but near-zero weights for the GRBS, and so on. To quantify number of voxels and their combined weights in each octant, we compute the sum of squared distances from the origin (0,0).

## Results

### Behavioral Results

Table S1 summaries the behavioral results of Study 1 (see also Yu et al. 2014). Essentially, participants felt highest level of guilt in the Pain: Self_Responsible condition, less so in the Pain: Both_Responsible condition and still less in the Pain: Partner_Responsible condition (*F* (2, 46) = 33.43, *p* < 0.001). This pattern was also observed for the amount of pain stimulation the participants chose to bear for the partner (*F* (2, 46) = 65.09, *p* < 0.001), and the perceived responsibility in causing the pain stimulation (*F* (2, 46) = 35.31, *p* < 0.001). Post-hoc tests showed that all comparisons between conditions exhibited significant difference for all the three measures (*p*s < 0.007).

**Table 1.**
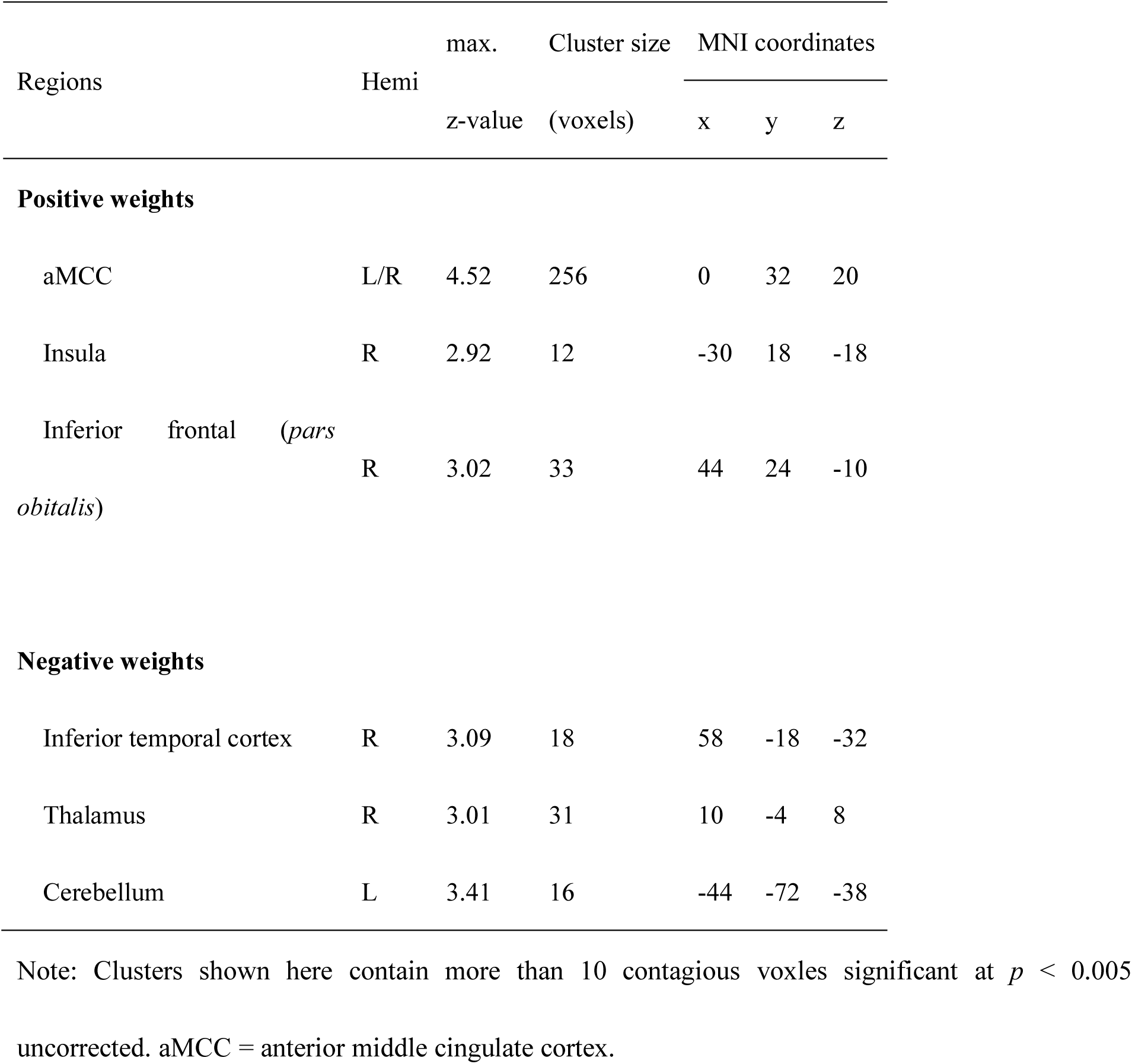
Activations in the thresholded GRBS map

Table S2 summaries the behavioral results of Study 2 (see also Koban et al. 2013). Post-scan self-reported guilt, but not other emotions, was higher for the “Play: Error_Pain” condition than the “Observe: Error_Pain” condition (Pairwise Bonferroni-corrected comparisons with sign tests, *Z* = 2.9, *p* = 0.003). In particular, the emotion shame, which frequently co-occur and is easily confused with guilt in everyday usage (Boonin, 1983; Fessler, 2004), showed a dissociable pattern in response to our manipulation. Specifically, self-reported guilt was significantly higher than self-reported shame in the “Play: Error_Pain” condition (mean difference 0.90 ± 0.40, *p* = 0.037, Bonferroni-corrected for multiple comparisons), but not in the “Observe: Error_Pain” condition (mean difference 0.05±0.05, *p* = 0.331), as supported by a significant Emotion type (guilt vs. shame) by condition (“Play: Error_Pain” vs. “Observe: Error_Pain”) interaction (*F*(1, 18) = 4.45, *p* = 0.049). Taken together, the self-reports results confirmed our hypothesis that one’s own responsibility in causing harm to others is a crucial cognitive process (or antecedent) underlying guilt. Further details regarding behavioral results can be found in Yu et al. (2014) and Koban et al. (2013).

### Neuroimaging results

#### Testing the sensitivity and generalizability of the Guilt-related Brain Signature (GRBS)

To determine whether there is a multivariate pattern that can distinguish between the social situation where the participants were solely responsible for others’ harm (i.e., high guilt state) and the situation where they were less causally responsible (i.e., low guilt state), we trained a linear support vector machine (SVM) to discriminate the “Pain: Self_Responsible” condition and the “Pain: Both_Responsible” condition with a leave-one-subject-out cross-validation (Friedman et al. 2001). The reason of choosing these two conditions for the comparison is that it rules out potential contamination by the feedback of participants’ own performance (i.e., correct vs. incorrect guess). Figure 2A shows the unthresholded GRBS weight map within the ‘Emotion’ meta-analytic map. As can be seen, the anterior middle cingulate cortex (aMCC), dorsomedial prefrontal cortex, bilateral insula, and the midbrain (including the periaqueductal grey, PAG) exhibited high positive predictive weights for detecting a guilt state (Table 1). For illustration purpose, we show a thresholded weight map obtained from a bootstrap procedure (5000 iterations, *z* > 2; Fig. 2A inset). It should be noted that the weight map is a distributed pattern in which all the voxels in the Emotion mask contribute to the classification. Examples of unthresholded patterns within aMCC and right AI are presented in the insets.

**Figure 2.**
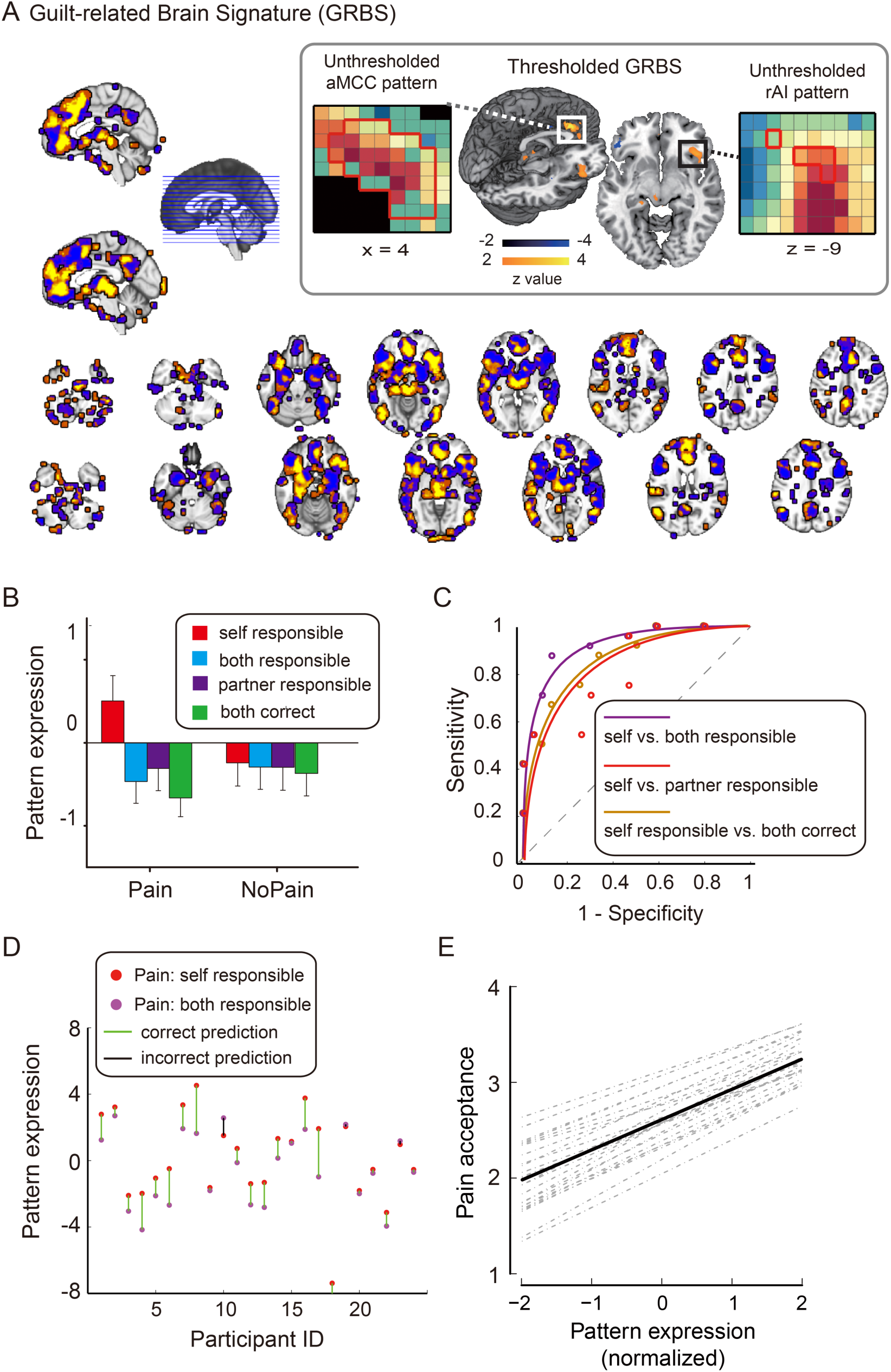
Guilt-related Brain Signature (GRBS) and its sensitivity. (A) Between-participant SVM weight map for guilt states (unthresholded). Bootstrap thresholded maps (5000 interations, *z* > 2) is shown in the inset. Examples of unthresholded patterns within right insula (rAI) and anterior middle cingulate cortex (aMCC) are also presented in the inset; small colored squares indicate voxel weights, black squares indicates empty voxels located outside of the GRBS pattern, and red-outlined squares indicate significance at *p* < 0.005 uncorrected (see also Table 1); (B) Cross-validated pattern expression computed as the dot product of the GRBS with the activation contrast maps for each participant; (C) Receiver operating characteristic curves (ROC) for the two-choice forced-alternative accuracies for the training dataset (Study 1). Purple: “Pain: Self_Responsible” vs. “Pain: Both_Responsible”; Red: “Pain_Self_Responsible” vs. “Pain: Partner_Responsible”; Gold: “Pain: Self_Responsible” vs. “Both_Correct”. (D) Individual participants’ pattern expression values for the self_err and both_err conditions. (E) The pattern expression values in the three errorous conditions in the Pain block (i.e., Self_Responsible, Both_Responsible, and Partner_Responsible) were predictive of the amount of pain sitmulation the participants were willing to tolerate for the confederates.

Pattern expression values reflect the distance between a given activation map and the classifier represented by a hyperplane represented in the feature space. To obtain single pattern expression values for each condition and each participant, we computed the dot product of the cross-validated weight map of the guilt pattern and the individual contrast images. These pattern expression values were then tested for differences between experimental conditions (Fig. 2B). We computed the forced-choice classification accuracy for how well the two conditions in questions were correctly classified based on their pattern expression values. Receiver-Operating-Characteristic (ROC) curve was created based on the performance of the classification. Pattern expression of GRBS for the eight conditions in Study 1 showed a significant Block (Pain vs. NoPain) by Outcome (Self_Responsible, Both_Responsible, Partner_Responsible, and Both_Correct) interaction, *F* (3, 69) = 7.68, *p* < 0.001. Planned comparisons showed that the pattern expression for the Pain: Self_Responsible was significantly higher than all the other three conditions in the Pain block (*p*s < 0.05; Fig. 2C), while the pattern expression of the other three Pain conditions did not differ significantly between one another. As shown by the ROC curves in Figure 2C, the GRBS discriminated “Pain: Self Responsible” versus “Pain: Both Responsible” with 88% (± 7%) accuracy (*p* < 0.001), “Pain: Self Responsible” versus “Pain: Partner Responsible” with 71% (±10%) accuracy (*p* = 0.064), and “Pain: Self Responsible” versus “Pain: Both Correct” with 75% (±11%) accuracy (*p* = 0.023). For the NoPain block, the only significant difference in the pairwise comparison was between NoPain: Self_Responsible and NoPain: Both_Correct (*p* = 0.021).

The majority of the participants (21 out of 24) exhibited larger pattern expression for the Self_Responsible than for the Both_Responsible conditions (Pain block; Fig. 2D). Moreover, regression analysis showed that the pattern expression values in the Pain block was predictive of pain sharing choice (i.e., reparation) (*b*_pattern_ = 0.092±0.036, *t* = 2.55, *p* = 0.015), suggesting that the GRBS contains information for atonement in guilt states (Fig. 2E).

We then tested whether the predictive power of the GRBS can be generalized to Study 2, another fMRI dataset using a similar interpersonal transgression paradigm (Koban et al. 2013). To this end, we computed pattern expressions of the guilt pattern applied to each condition of Study 2. Pattern expression of the GRBS for the six conditions showed a significant Feedback-by-Agency interaction, *F* (2, 36) = 4.59, *p* = 0.013 (Fig. 3A). Pairwise comparisons showed that the pattern expression for the Play: Error_Pain was significantly higher than the Play: Correct (*p* = 0.002), the Observe: Error_Pain (*p* = 0.016), and marginally significantly higher than the Play: Error_Warmth condition (*p* = 0.075). Pattern expression of the Observe conditions did not differ significantly between one another (Fig. 3A). We then tested the classification accuracy based on these pattern expression values. As shown in Figure 3B, the GRBS discriminated “Play: Error_Pain” versus “Observe: Error_Pain” with 74% (±10%) accuracy (*p* = 0.064), “Play: Error_Pain” versus “Play: Error_Warmth” with 74% (±10%) accuracy (*p* = 0.064), “Play: Error_Pain” versus “Play: Correct” with 79% (±9%) accuracy (*p* = 0.019), and “Play: Error_Pain” versus “Observe: Correct” with 79% (±9%) accuracy (*p* = 0.019). In sum, these results show that the predictive power of GRBP generalizes to a novel and completely independent dataset of interpersonal guilt.

**Figure 3.**
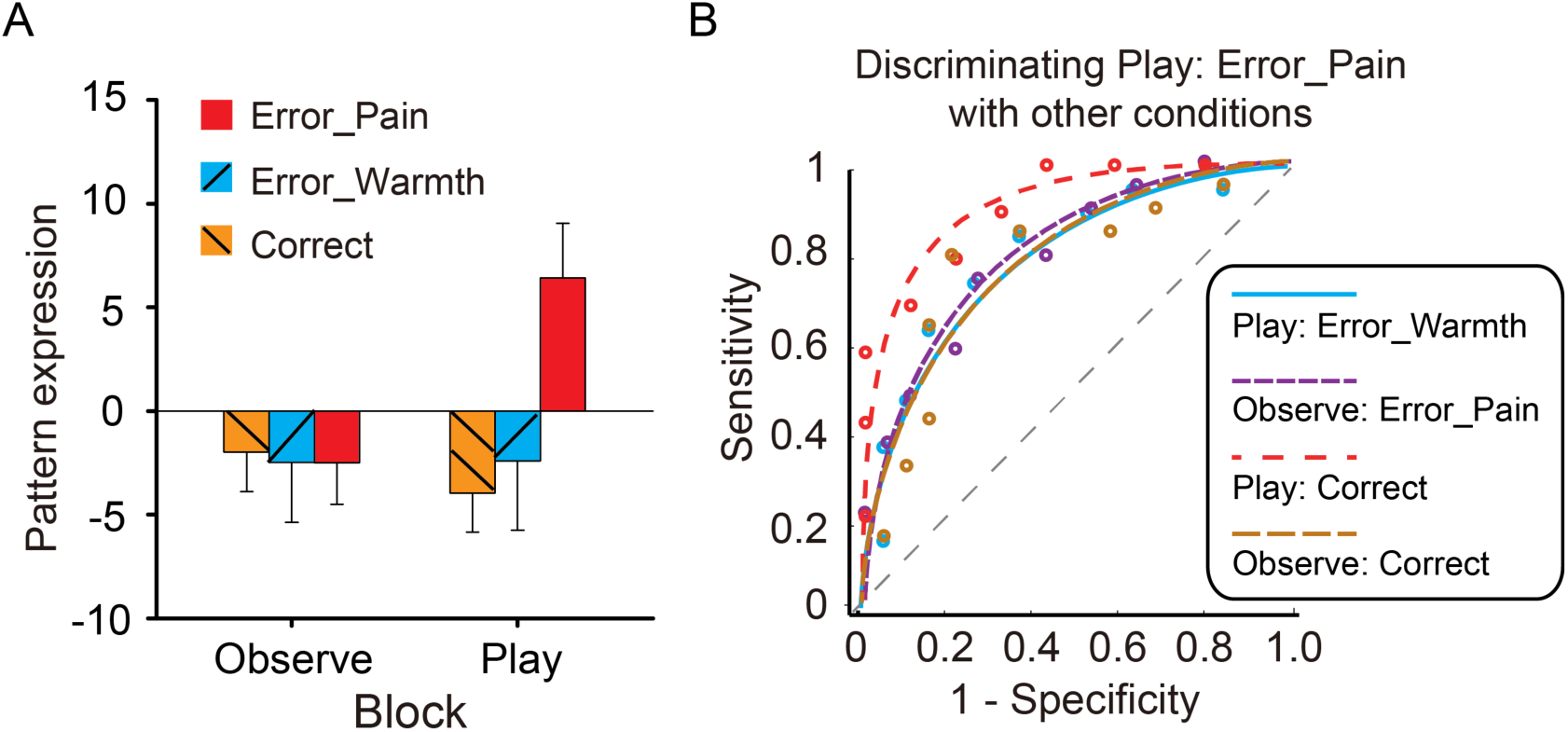
Generalization of the GRBS. (A) In Study 2 dataset, the “Play: Error_Pain” condition (i.e., the condition associated with highest guilt) shows the highest pattern expression. In this condition, the participant’s action caused pain. In “Warmth” conditions, the participant’s action did not cause pain to the partner. In “Correct” conditions, the participant did not make an error and no punishment was delivered to the partner. In “Observe” conditions, the participant observed the game and pain was not contingent on their actions. (B) ROC curves for the two-choice forced-alternative performance for the validation dataset (Study 2). Blue: Play Error Pain vs. Play Error Warmth; Purple: Play Error Pain vs. Observe Error Pain; Red: Play Error Pain vs. Play Correct; Gold: Play Error Pain vs. Observe Correct. Error bars indicate s.e.m.

In Study 1 and Study 2, we manipulated two critical antecedents of guilt interpersonal harm and one’s own responsibility in causing that harm (Koban et al. 2013; Yu et al. 2014). The guilt signature therefore should capture their super-additive interaction. The performance of our guilt signature met this criterion: it did not discriminate levels of responsibility in causing non-harmful consequences (e.g., Study 1, NoPain: Self Responsible vs. NoPain: Both Responsible conditions, accuracy = 58%±10%, *p* = 0.54), nor did it discriminate harmful from non-harmful consequences for which the participants were not responsible (e.g., Study 2, Observe: Error_Pain vs. Observe: Error_Warmth, accuracy = 47%±12%, *p* = 1; Observe: Error_Pain vs. Observe: Correct, accuracy = 53%±12%, *p* = 1). However, it did respond, as we showed above, when participants were responsible and causing harm.

#### Testing the specificity of the GRBS

To assess the specificity of the classifier, we examined its predictive power in two other independent data sets: one using thermal (heat) pain and observed (vicarious) pain (Krishnan et al. 2016), the other using recall task to elicit basic and social emotions (Wagner et al. 2011). Univariate analyses reported in these previous studies have implicated the brain regions showing highest predictive weights in the GRBS (e.g., aMCC, rAI) in the processing of physical and vicarious pain, and in the processing of recalled guilt episodes. However, it is an open question whether these brain states are distinguishable to GRBS. The multivariate approach allows us to test whether shared univariate activations reflect common neural representations (Woo et al. 2014). As can be seen from Figure 4 (see also Table S3), GRBS performed at chance level in discriminating different intensity of thermal pain stimulation (High vs. Medium: accuracy = 57±11%, *p* = 0.57; Medium vs. Low: accuracy = 46±9%, *p* = 0.85) and different degree of vicarious pain (High vs. Medium: accuracy = 50±9%, *p* > 0.99; Medium vs. Low: accuracy = 57±9%, *p* = 0.57). The classifier did not significantly differentiate recalled guilt from either recalled sad memories (accuracy = 33±12%, *p* = 0.30) or recalled shame memories (accuracy = 60± 13%, *p* = 0.61). These findings suggest that GRBS is better at detecting transgression in real-time interpersonal contexts than other unpleasant experiences, including guilt-related memories. That is, it does not appear to be selectively activated during retrieval of guilt-related memories, but it does respond selectively to feedback indicating that one has caused harm to a partner and predicts atonement behavior.

**Figure 4.**
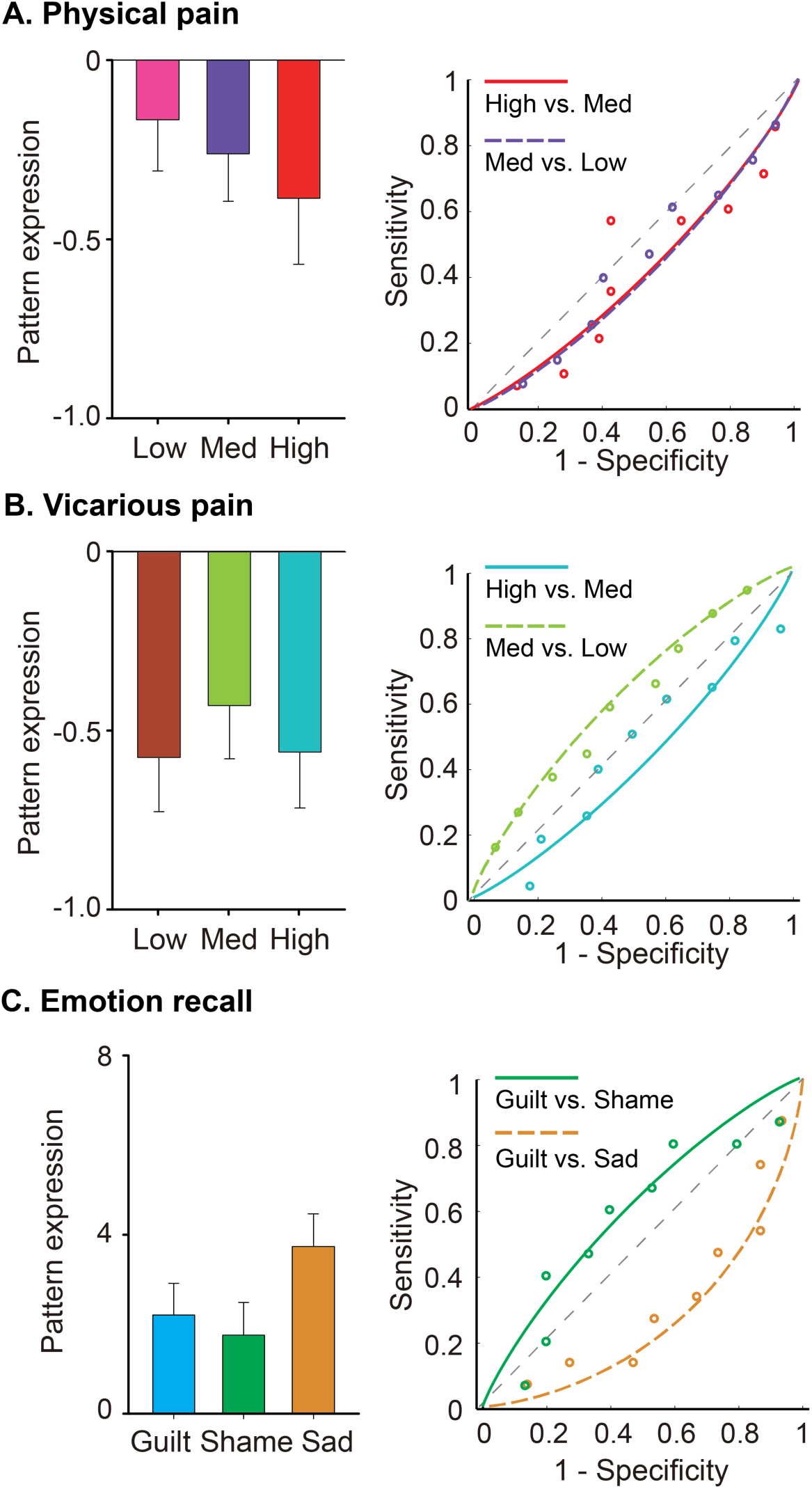
Specificity of the GRBS. (A-C) Pattern expression and ROC curves for a thermal pain dataset (A), a vicarous pain dataset (B) and an emotion recall dataset (C). GRBS cannot discriminate different levels of physical pain, vicarous pain, or different types of emotional memories (including guilt-related memories), suggesting that the predictive power of GRBS was specific to detecting one’s transgression in the immediate social interaction context (see also Table S3). Error bars indicate s.e.m.

Finally, we investigated the relationship of the GRBS to other, potentially similar brain signatures of social-affective processes. Spatial similarity (Pearson correlation coefficients across all voxels) between the GRBS and eight other brain signatures are shown in Table S4 and Figure S2. Most patterns showed around zero correlation (*r*’s between −0.1 and 0.1), with the exception of the PINES—developed to track negative affect associated with unpleasant images (Chang, et al. 2015)—, which showed a weak positive correlation (*r* = 0.12) with GRBS, thus suggesting some shared variance between those two brain patterns. To examine this similarity more closely, we qualitatively examined whether it might be driven by shared positive or negative weights in ACC or insula, or other areas often activated by emotional events, such as the amygdala (ROIs defined based on anatomical labels and the WFU Pickatlas version 3.0.5b (Maldjian et al. 2003)). Figure 5A shows the joint distribution of normalized (*z*-scored) voxel weights of PINES on the x-axis and GRBS on the y-axis (cf. Koban et al. 2019). Differently colored octants indicate voxels of shared positive or shared negative (Octants 2 and 6, respectively), selectively positive weights for GRBS (Octant 1) and for PINES (Octant 3), selectively negative weights for GRBS (Octant 5) and for PINES (Octant 7), and voxels where the voxel weights of the two signatures went in opposite directions (Octants 4 and 8) (Fig. 5B). Overall correlations between the two patterns in the emotion mask (Fig. 5A) and in the three ROIs (Fig. 5C-5E) were relatively weak. Across the whole emotion mask, stronger weights (sum of squared distances to the origin [SSDO]) were actually observed in the non-shared octants (1,3,5,7). Further, the three ROIs showed distinct patterns of covariation between the two patterns. Many voxels in the bilateral amygdalae showed positive weights for PINES, but not for GRBS, as reflected by the high SSDO in Octant 3 (Fig. 5C). This is in line with the long-established role of the amygdala in emotional attention (see Vuilleumier, 2005 for a review) and in assigning affective salience to sensory stimuli (LeDoux, 2000). Bilateral insulae showed strongest weights in the Octants 1,2, and 7, indicating many positive weights for guilt specifically (Octant1), as well as shared positive weights across the two signatures (Octant 2), but also some many voxels with negative weights in the PINES (Octants 6-8) (Fig. 5D). Finally, the ACC showed almost exclusively positive weights for GRBS, which were mostly near-zero or even negative for PINES (Octants 1 and 8) (Fig. 5E). Thus, while the insula might include some shared positive weights, the overall results suggest distinct activation patterns for guilt and picture-induced negative affect in emotion-related brain areas.

**Figure 5.**
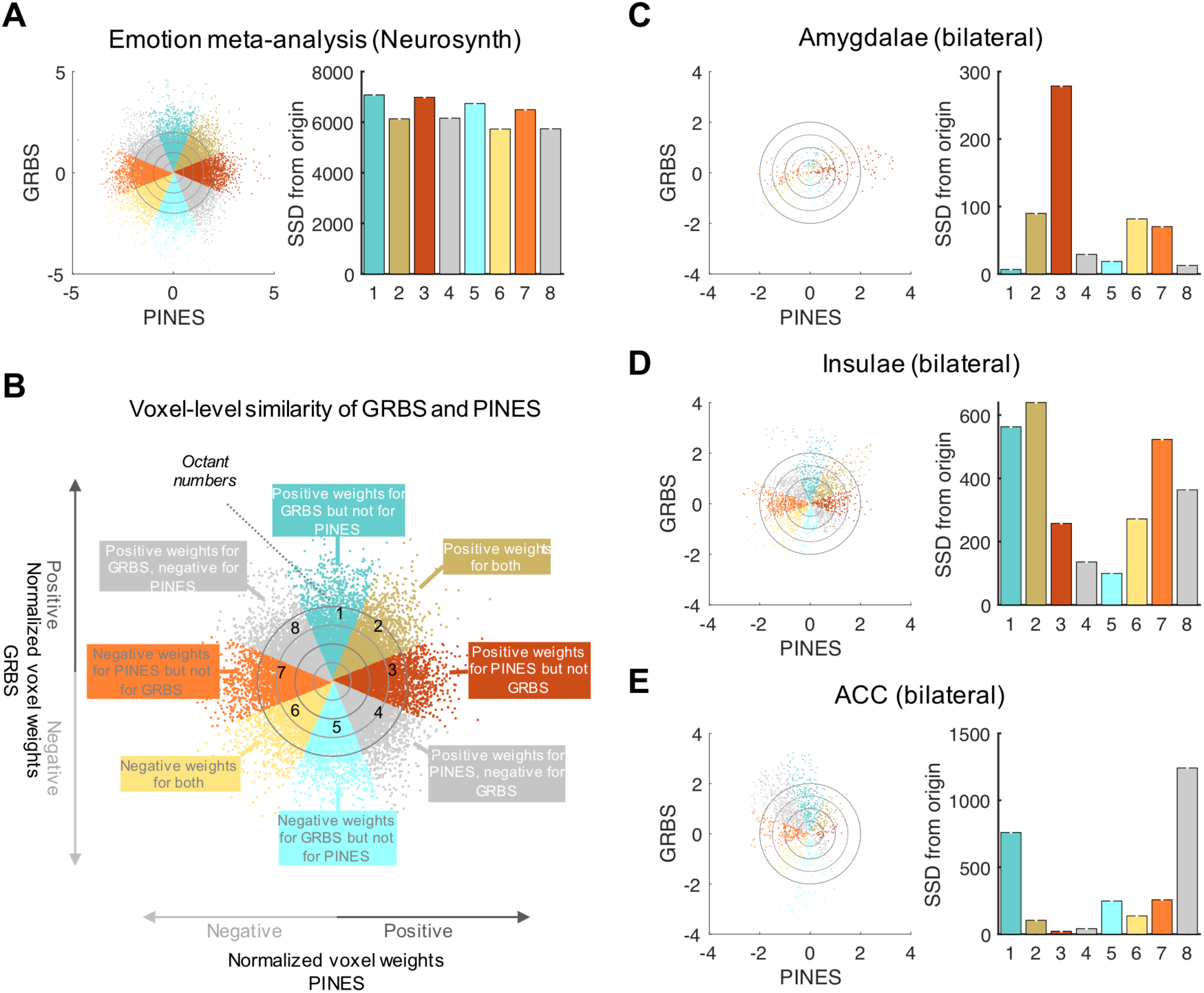
Voxel-level spatial similarity between GRBS and PINES. (A) Scatter plots displays normalized voxel (within the Emotion mask) beta weights for GRBS (y-axis) and PINES (x-axis). Bars on the right represent the sum of squared distances from the origin (0,0) for each octant. This value integrates the number of voxels and their combined weights in each octant, we compute. (B) Differently colored octants indicate voxels of shared positive or shared negative (Octants 2 and 6, respectively), selectively positive weights for GRBS (Octant 1) and for PINES (Octant 3), selectively negative weights for GRBS (Octant 5) and for PINES (Octant 7), and voxels where the voxel weights of the two signatures went in opposite directions (Octants 4 and 8). (C) Voxel-level spatial similarity in bilateral amygdalae shows positive weights for PINES, but not for GRBS, as reflected by the high SSDO in Octant 3. (D) Voxel-level spatial similarity in bilateral insulae shows strongest weights in the Octants 1, 2 and 7, indicating many positive weights for guilt specifically (Octant1), as well as shared positive weights across the two signatures (Octant 2), but also some many voxels with negative weights in the PINES (Octants 6-8). (E) Voxel-level spatial similarity in ACC shows almost exclusively positive weights for GRBS, which were mostly near-zero or even negative for PINES (Octants 1 and 8).

## Discussion

Characterizing how specific emotions are generated and represented in the brain is a central question in affective neuroscience and important for understanding emotions and their regulation in healthy and clinical individuals (Hamann 2012; Bijsterbosch et al. 2018). However, given the substantial overlap between brain correlates of different psychological processes, including positive and negative emotions (Kober et al. 2008; Lindquist and Barrett 2012; Wager et al. 2015), identifying distinct brain correlates of different emotions has proven to be a very challenging goal, which may require multivariate approaches that go beyond contributions of single brain regions (Woo et al. 2014; Kragel and LaBar 2015; Skerry and Saxe 2015; Wager et al. 2015). The present results contribute to this undertaking by providing first evidence that even complex social or moral emotions such as guilt can be accurately identified based on a distributed multivariate brain pattern—the Guilt-related Brain Signature (GRBS). Developing a multivariate pattern for detecting the presence of guilt-related psychological states helps us understand the neural mechanism underlying guilt and atonement, and serves as a tool for future studies that aim at manipulating and/or measuring guilt in different environments and populations (Wager et al. 2013; Chang et al. 2015; Krishnan et al. 2016).

Interpersonal guilt reflects the ability to detect and respond to a situation where someone else is harmed and in which oneself is the source of that harm (Boonin, 1983). This type of guilt is thought to be critical for maintaining social norms and interpersonal relationships (Baumeister et al. 1994). On the transgressor’s side, accurately detecting such a situation and reacting appropriately allows them to restore the reputation and social relationship with the victim (via direct reciprocity; Yu et al. 2014) and other relevant individuals in the social network (via indirect reciprocity and social image; Stearns and Parrott 2012). Moreover, the transgressor’s expression of guilt and conciliatory gestures reaffirm the abiding power of the violated social norms and compensate the loss of the victim (Bicchieri 2005).

Paralleling an approach used for other social-affective processes (Woo et al. 2014; Krishnan et al. 2016), we used SVM on fMRI data to classify the presence versus absence of the core appraisal of guilt, namely, one’s responsibility in causing harm to another. The GRBS had good cross-validated predictive accuracy (71% – 88%) and significantly predicted compensation behavior, thus linking cognitive brain processes to relevant behavioral outcomes. Further, the GRBS showed high accuracy (74% – 79%) on a completely independent test set from a different laboratory and culture, demonstrating its robustness to variations in experimental settings and cultural context.

This signature, while being a distributed pattern across the entire “Emotion” network (Yarkoni et al. 2011), exhibits its highest predictive weight in the aMCC and right anterior insula (Fig. 2A). Although these peak voxels parallel the previous univariate analyses (Koban et al. 2013; Fourie et al. 2014; Yu et al. 2014; Cui et al. 2015), they nevertheless contribute independently to the understanding of the neurocognitive mechanism of detecting one’s transgression and reacting accordingly. The multivariate analysis derives a weight map that captures the core processes underlying interpersonal transgression and guilt. This abstract weight map can then be applied to new observations from the same or different datasets to assess its sensitivity, specificity, and generalization (Wager et al. 2013). Specifically, when it comes to the aMCC and anterior insula, extensive research, including those of our own, has demonstrated the lack of functional specificity in these areas using neuroimaging meta-analyses and multivariate pattern analysis (e.g., Lindquist et al. 2012; Wager et al. 2016; Yarkoni et al. 2011). We have also argued, and provided evidence, that multivariate pattern-related activity in such areas offers greater functional specificity than simply interpreting overlapping activation (Kragel et al. 2018). For example, in Kragel et al. (2018), we found that the aMCC contains a population-level multivariate representation (pattern) related to pain that generalizes across 3 types of somatic pain (tested across 6 studies), but is not shared by 3 kinds of negative emotion tasks or 3 kinds of cognitive control tasks. We argue that multivariate pattern analysis works because it picks up, to some degree, on differential patterns of activation across neural populations (and microvasculature) that are unevenly distributed across voxels (for review and discussion, see Kragel et al. 2018).

Supporting the notion of distinct multivariate patterns for different affective processes, we found only weak correlations between the GRBS and other pain- and emotion-related brain patterns, such as the PINES. Further, even local patterns in emotion-related areas—including the ACC and insula—showed only limited shared variance between the GRBS and the PINES. Interestingly, the patterns of shared versus unique weights for the two signatures were distinct across the three regions of interest. The insula showed some evidence of common positive weights for both GRBS and PINES, which is in line with partially shared processes. In contrast, amygdala and ACC voxels with positive weights for one signature were often near-zero or had even negative weights in the other signature, suggesting very distinct local contributions to the overall patterns.

Moreover, in the current study, the signature was derived from a sample of Chinese participants (East Asian culture) and the predictive power of this pattern can be partially generalized to a sample of Caucasian participants (Western culture), suggesting that the core underlying neurocognitive processes may be similar even across different cultures and experimental settings. The GRBS was also sensitive to the levels of guilt (as manipulated via agency of another’s pain) in the interactive action monitoring task. Yet, the signature did not discriminate levels of either physical pain (i.e., receiving painful stimulation; Krishnan et al. 2016) or vicarious pain (i.e., observing others receiving painful stimulation; Krishnan et al. 2016; Fig. 3A-3C), which are both arousing, aversive, salient experiences. Interestingly, the signature did not discriminate guilt-related memories from other type of negative emotional memory either (Fig. 3D-3F). Memories of guilt episodes may involve recognition of one’s causality in other’s suffering, but likely do not involve the processes of detecting and responding to such components in the here-and-now social context (Redcay and Schilbach, 2019). Taken together, our findings suggest that in an interpersonal transgression context, the transgressor’s brain does not only capture the distressful consequence of others *per se*, as in the case of experiencing vicarious pain, but also actively seeks the attribution of the harm and, when one’s own responsibility is confirmed, decides how to respond (e.g., atonement, apology). This finding, together with the predictive power of the GBRS in tracking reparation behavior (i.e., compensation), suggests that brain activation patterns identified here may primarily implicate the impact of guilt-related appraisal on subsequent behavioral responses, in line with the notion that emotions serve to guide adaptive behaviors and generate corresponding action tendencies. These effects may be absent in recalled guilt, thus precluding a successful decoding of GBRS in this condition

Further, we note that individual differences in GRBS responses were not predictive of guilt ratings in either dataset. One explanation is that the ratings of subjective feelings of guilt were collected after the task in the scanner and thus were simply recall in nature, whereas the GRBS, as our results show, is specific to detecting and responding to immediate transgression. Alternatively, the individual differences in GRBS response may be influenced by other factors such as overall signal, and our sample may be underpowered to detect small between-person correlations. Future studies that simultaneously record fMRI and more sensitive online measures of emotional feeling of guilt (e.g., eye gaze pattern, skin conductance; see Yu et al. 2017) may be able to explore GRBS’s roles in the temporal unfolding of guilt experience. Namely detecting the presence of cognitive antecedents of guilt, encoding guilt feelings as experienced immediately in interpersonal transgression, and predicting atonement following guilt (Amodio et al. 2007). More broadly, the multivariate approach can inform our understanding of the neural basis of social cognition by developing brain signatures that capture specifically defined cognitive processes and testing their generalizability to other social cognitive functions. This way, we would be able to restructure our understanding of social cognition on the basis of underlying brain representations.

A conceptual clarification about guilt and responsibility is worth noting. In this paper, “guilt” refers to a constellation of cognitive-affective processes in response to interpersonal transgression and harm (e.g., detecting harm and assigning responsibility), rather than simply the feeling/experiential component of this constellation of processes. On this conceptualization of guilt, recognizing one’s causal responsibility is an integral part of guilt (Ellsworth and Smith 1988; Tracy and Robins 2006), rather than an independent process that is parallel to guilt, at least in most situations. Nevertheless, we acknowledge that it is an interesting and important empirical question as to whether guilt feelings can arise, in certain populations or circumstances, without objective causal responsibility in interpersonal harm. For example, survivors of disasters or atrocities sometime report that they experience “guilty” feelings towards other victims who suffer much more than they do, despite the fact that they are not causally responsible for other victims’ suffering. One possible psychological mechanism underlying such “survivor guilt” is that survivors falsely attribute responsibility of others’ suffering to themselves (O’Connor et al. 2000). Similarly, “existential” guilt, negative feelings towards oneself as a purposeless or unworthy being experienced by people with certain type of depression, seems to be a result of illusory perceptions of responsibility (Ratcliffe, 2014). Conversely, some individuals (e.g., those high in psychopathy; Cima et al. 2010) may have the attribution of responsibility for harm without feeling guilt. The guilt signature could be used as a tool to empirically test these hypotheses. Unfortunately, direct tests of these interesting possibilities are beyond the scope of this paper, and await further studies designed for this purpose.

It may be argued that the term ‘guilt’ is not used equivalently across Chinese and Swiss cultures and languages. This is related to a more profound issue as to how we could know whether or not people living in different cultures and speaking different languages are experiencing the same *kind* of emotion when they claim that they are feeling guilty (English), or schuldig (German), or coupable (French), or nei jiu (Chinese)? In this study, we adopt the assumption that ‘guilt’ refers to a category of emotional states, under which different variants of guilt are species with variant-specific defining features or differentia. The specific type of guilt that we investigated in this study, as we have argued, is defined by two critical features (Baumeister et al. 1994; Tracy and Robins, 2006): (1) recognizing a breach of moral norms, typically involving harm to another, and (2) attributing causal responsibility in such violation to oneself. These two features have been demonstrated to be reliable cognitive antecedents of guilt in both Western and East Asian cultures (Benedict, 1946/2005; Piers and Singer, 1971; Bedford and Hwang, 2003; Wong and Tsai, 2007), and have been manipulated to induce guilt, in both Western (Bastin et al. 2016; Cracco et al. 2015; Koban et al. 2013; Seara-Cardoso et al. 2016) and East Asian participants (Leng et al. 2017; Furukawa et al. 2019; Yu et al. 2014; Zhu et al. 2019). In line with these theoretical and empirical work, we utilized these two defining features of guilt in the tasks of our training and test datasets. Importantly, it is not required that participants from all cultures experience this type of guilt to the same degree in response to the same situations. Our analyses require only that it is experienced to some degree by participants across cultures. We showed that the guilt-related pattern we identified was indeed preserved cross-culturally, at least in the context of our study, thereby providing empirical support for common cross-cultural brain processes. This finding extends the commonality in the cognitive-affective processes underlying guilt to the level of (partially) shared cognitive-affective processes underlying guilt and its brain correlates across cultures and context. It is an interesting and important empirical question for future research as to what extent this signature could discriminate different variants of guilt both within and across cultures (i.e., causing physical harm versus social harm; causing harm to a friend versus a stranger).

To be sure, we are not the first to explore how emotions arise by relating appraisal theory with pattern recognition analyses of human neuroimaging data (for a review, see Adolphs, 2017). For example, Skerry and Saxe (2015) show that discrete emotion categories that people assign to a given emotion-eliciting event can be accurately predicted by a set of abstract features of the events (e.g., whether the protagonist is responsible for the outcome in the event). This abstract feature-based model outperformances the predictions based on two other influential models of emotion (i.e., the basic emotion theory and the arousal-valence theory). Adopting a similar theoretical framework (i.e., the appraisal theory of emotion), our study can be seen as a case study focusing on interpersonal guilt, with responsibility for harm to another as its core appraisal. In fact, in Skerry and Saxe (2015)’s fine-grained feature space consisted of 38 appraisal dimensions, the feature “caused by self” is most consistently highlighted to be relevant to guilt. Future research could leverage this feature-space approach to formally test psychologically meaningful hypotheses concerning the distinction (or the lack thereof) between guilt and other related social and non-social emotions, such as shame, embarrassment, and non-social regret. This approach also provides an interesting, brain-based way to compare emotions across cultures. Although a one-to-one mapping of emotion terms across languages may be problematic, abstract event features may be less likely to be ‘lost in translation’ and shared cross-culturally (Hurtado de Mendoza et al. 2010; Fiske, 2019).

The generalizability of the GRBS to Study 2 seems limited. In particular, the difference between the pattern expression of the “Play: Error_Pain” condition and that of the “Play: Error_Warmth” condition was at trend level. This might be in part due to the small sample size of Study 2. Another conjecture is that given that the classifier was trained to discriminate one’s causal responsibility in interpersonal harm, it might be more sensitive in detecting differences in responsibility than in detecting differences in the severity of harm. This is supported by the fact that the signature responded more distinctively to “Play: Error_Pain” versus “Observe: Error_Pain”, two conditions that differ only in appraisals of responsibility but not in severity of harm. Therefore, we acknowledge that our goal of developing a sensitive, specific and generalizable brain signature of guilt has yet to be fully achieved; but we believe the present multivariate pattern is both useful and a critical motivating stepping stone to large-scale studies that would be required to perform a more definitive identification of cross-cultural neural representations of guilt. Future studies are needed to achieve this goal in larger samples and to incorporate more fine-grained manipulation of guilt (e.g., responsibility, severity of harm, relationship between transgressors and victims, etc.). Nevertheless, the utility of a provisional model such as ours might become clearer when compared with measures in other domains that have been only partially validated and/or have limited specificity. For example, face-related activity is routinely identified in fMRI in individuals, and in spite of little to no validation of its specificity to faces in that individual (and debatable specificity of the general area of the ‘fusiform face area’), it is routinely used to infer the persistence of face-related representations in working memory (Druzgal and D’Esposito 2001; Lewis-Peacock and Postle 2008), long-term memory (Polyn et al. 2005), attention (Yeung et al. 2006), and others. Even relative or limited specificity is reasonable for use of an fMRI pattern for further testing, though the wisdom of doing so must be evaluated on a case by case basis. In a similar vein, there are very few biomarkers in medicine that are highly sensitive and specific, but even moderate diagnostic value confers information. In the same spirit, our guilt-related pattern confers information value, as well as a defined brain measure, for provisional inference, brain comparisons, and further testing and validation on the brain bases of social emotions.

To conclude, we developed a neural signature, the Guilt-related Brain Signature (GRBS), that is sensitive and specific to the critical appraisals underlying the experience of guilt in social interactions, namely, recognizing one’s responsibility in causing other’s suffering (Frijda 1993; Baumeister et al. 1994). Showing its predictive validity for behavioral outcomes, the response of this signature predicts atonement decisions following transgression even after statistically controlling for experimentally manipulated degree of responsibility. Supporting its discriminative validity, the GRBS did not respond to guilt memories or memories of other negative emotions, neither did GRBS respond differently to increasing levels of vicarious pain or increasing levels of agency in non-harmful outcomes. It was also not strongly correlated with any other previously developed affect- or pain-related signature, ruling out the possibility that it reflects general negative affect or other related categories of social emotions like empathy for pain and perception of self-agency. This signature can be used in future studies for detecting guilt- and transgression-related neural processes, for example by manipulating other important social factors, such as intentions of transgression and interpersonal relationship between transgressors and victims, by applying it to harm-based moral decision-making context (Yu, Siegel, Crockett, 2019), or by testing its response in different clinical populations such as those characterized by excessive or reduced experience of guilt (i.e., internalizing disorders versus psychopathy).

## Acknowledgements

This work was supported by the following sources of funding: National Natural Science Foundation of China (31630034, 71942001) and National Basic Research Program of China (973 Program: 2015CB856400) awarded to XZ, the Royal Society Newton International Fellowship (NF160700) awarded to HY, NIH National Institute of Mental Health grant (R01 MH116026) awarded to LC and TW, NIH National Institute on Drug Abuse grant (R01 DA035484) awarded to TW, Swiss National Science Foundation (32003B_138413) and NCCR Affective Sciences at University of Geneva (51NF40-104897) awarded to PV. The authors thank Dr. Molly Crockett, Dr. Sheng Li, Dr. Jian Li, Dr. Miguel Eckstein, Dr. Philip Blue, Ms. Sophie Harrington and two anonymous reviewers for their helpful suggestions on data analysis and constructive comments on an earlier version of the manuscript.

## Supplementary information for

**Table S1.** Emotion Ratings of Study 1 (Means and SDs)

| Item | Pain block |  |  | *F* (2, 46) |
| --- | --- | --- | --- | --- |
|  | Partner_Responsible | Both_Responsible | Self_Responsible |  |
| Pain shared | 2.0 (0.6)a | 2.8 (0.6)b | 3.1 (0.6)c | 65.09*** |
| Responsibility | 3.0 (1.8)a | 4.6 (1.6)b | 7.1 (1.6)c | 35.31*** |
| Guilt | 1.8 (0.9)a | 3.4 (1.7)b | 5.3 (2.3)c | 33.43*** |
| Distress | 2.0 (1.5)a | 2.8 (2.0)a | 4.0 (2.4)b | 10.51*** |
| Fear | 2.6 (1.8) | 3.3 (2.3) | 2.9 (1.9) | 1.47 |
| Anger | 2.0 (1.6) | 2.6 (2.3) | 2.0 (1.3) | 1.89 |

Note: “Pain shared” is the average amount of pain the participants chose to take for the partner (1: “take none”; 4: “take all”). This measure was taken after each erroneous trial. “Responsibility” and emotions were rated on 1 – 9 scales after the MRI scanning. Standard deviations are shown in parentheses. Significant differences (critical $\alpha < 0.05$ Bonferroni corrected) in pair-wise comparison within each row are denoted by different subscripts. \*\*\* $P < 0.001$ in repeated measures ANOVA.

**Table S2.** Emotion Ratings of Study 2 (Means and SDs)

| Item | Play block |  |  | Observe block |  |  |
| --- | --- | --- | --- | --- | --- | --- |
|  | Correct | Error Warmth | Error Pain | Correct | Error Warmth | Error Pain |
| Fear | 1.2 (0.9) | 1.1 (0.3) | 1.9 (1.2) | 1.2 (0.9) | 1.4 (1.1) | 2.2 (1.4) |
| Shame | 1.0 (0.0) | 2.2 (1.4) | 3.2 (1.4) | 1.0 (0.0) | 1.1 (0.5) | 1.2 (0.9) |
| Guilt | 1.0 (0.0) | 2.5 (1.4) | 4.1 (1.1) | 1.0 (0.0) | 1.0 (0.0) | 1.3 (0.9) |
| Sadness | 1.0 (0.0) | 1.9 (1.5) | 2.2 (1.5) | 1.0 (0.0) | 1.3 (0.7) | 1.9 (1.5) |
| Anger | 1.0 (0.0) | 2.9 (1.4) | 3.3 (1.8) | 1.0 (0.0) | 1.1 (0.3) | 1.6 (1.0) |

Note: The five different emotions were rated on 1 – 5 Likert scales, after the main experimental blocks.

**Table S3.** Pattern expressions of GRBS in specificity tests.

| Item | High/Guilt | Medium/Shame | Low/Sad |
| --- | --- | --- | --- |
| Physical pain | -0.39 (0.98)* | -0.26 (0.70) | -0.17 (0.76) |
| Vicarious pain | -0.56 (0.83)** | -0.43 (0.79)** | -0.57 (0.80)*** |
| Emotion memory | 2.20 (2.73)** | 1.75 (2.81)* | 3.74 (2.82)*** |

*Note.* Standard errors are shown in parentheses. Significant differences relative to baseline (0) are denoted by asterisk: \* $p < 0.05$ , \*\* $p < 0.01$ , \*\*\* $p < 0.001$ . No significant difference was observed between conditions.

**Table S4.** Spatial similarity between GRBS and other social-affective brain signatures.

| Signature | Spatial similarity<br>(Pearson $r$) |
| --- | --- |
| NPS | -0.05 |
| PINES | 0.12 |
| Rejection | -0.02 |
| VPS | -0.04 |
| GSR | 0.09 |
| HR | -0.06 |
| Empathic Distress | -0.03 |
| Empathic Care | -0.03 |

*Note.* NPS = neurologic signature of physical pain (Wager et al., 2013), PINES = picture-induced negative emotion signature (Chang et al., 2015), Rejection = social rejection (Woo et al., 2014), VPS = vicarious pain signature (Krishnan et al., 2016), GSR = social threat-induced skin conductance signature (Eisenbarth et al., 2016), HR = social threat-induced heart rate signature (Eisenbarth et al., 2016), Empathic Distress = empathic distress signature (Ashar et al., 2017), Empathic Care = empathic care signature (Ashar et al., 2017).

**Figure S1.**
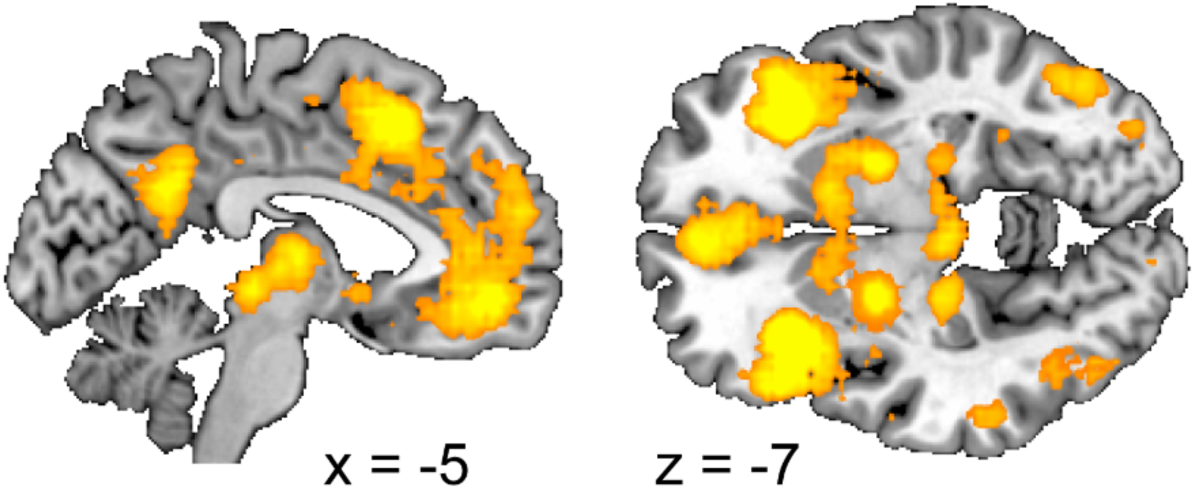
“Emotion” mask used in training the Guilt-related Brain Signature (GRBS). This is the uniformity test map (or the “forward inference map” in previous terminology) of the term “emotion” on the Neurosynth website (http://neurosynth.org/analyses/terms/emotion/, accessed on September 7th 2014).

**Figure S2.**
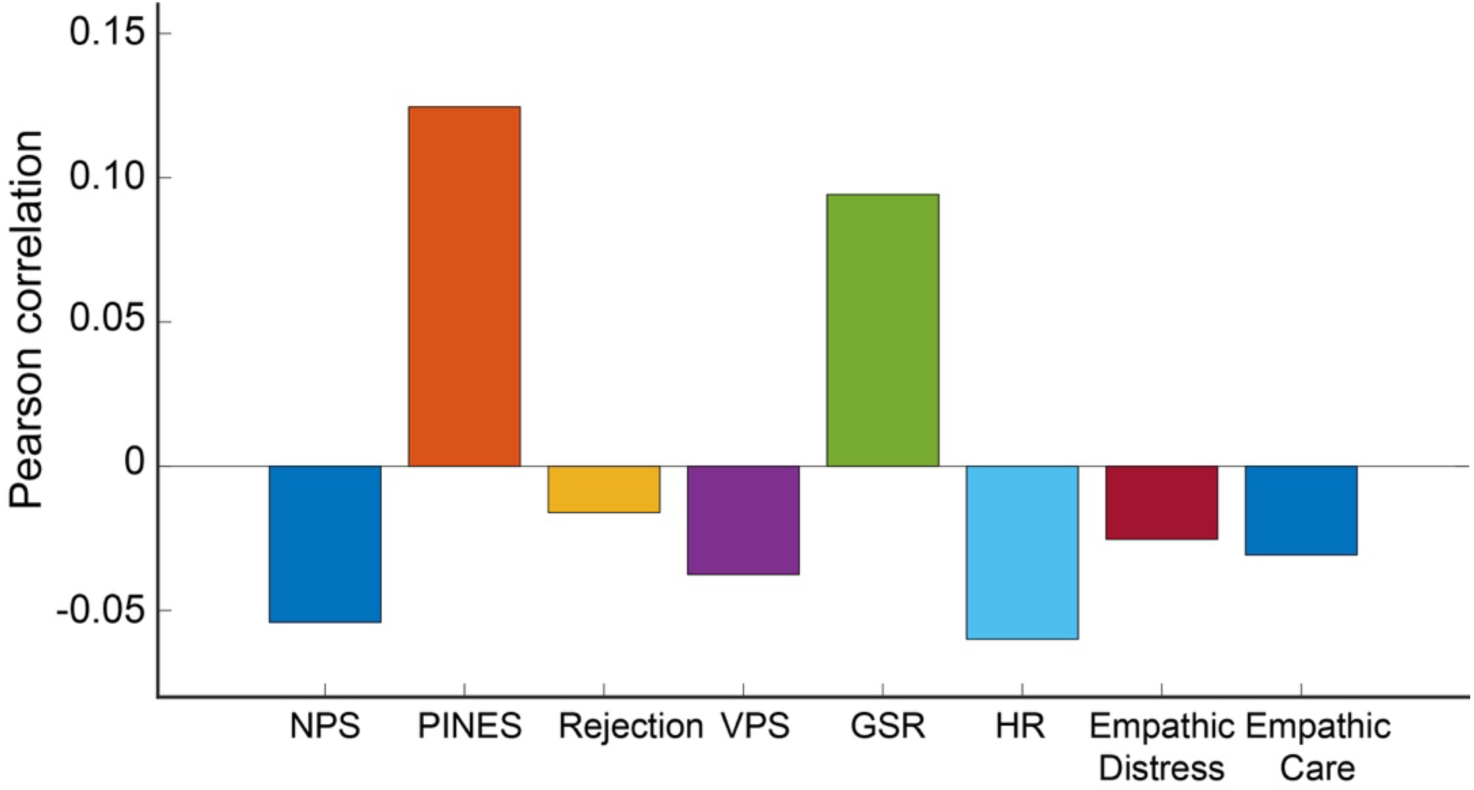
Spatial similarity (Pearson correlation) between the GRBS and eight other brain signatures. NPS = neurologic signature of physical pain (Wager et al., 2013), PINES = picture-induced negative emotion signature (Chang et al., 2015), Rejection = social rejection (Woo et al., 2014), VPS = vicarious pain signature (Krishnan et al., 2016), GSR = social threat-induced skin conductance signature (Eisenbarth et al., 2016), HR = social threat-induced heart rate signature (Eisenbarth et al., 2016), Empathic Distress = empathic distress signature (Ashar et al., 2017), Empathic Care = empathic care signature (Ashar et al., 2017).

## References

Adolphs, R. (2017). How should neuroscience study emotions? By distinguishing emotion states, concepts, and experiences. Social cognitive and affective neuroscience, 12(1), 24–31.

Amodio DM, Devine PG, Harmon-Jones E. 2007. A Dynamic model of guilt. Psychol Sci. 18:524–530.

Ashar YK, Andrews-Hanna JR, Dimidjian S, Wager TD. 2017. Empathic care and distress: Predictive brain markers and dissociable brain systems. Neuron. 94:1263–1273.

Barrett LF, Satpute AB. 2013. Large-scale brain networks in affective and social neuroscience: towards an integrative functional architecture of the brain. Curr Opin Neurobiol. 23:361–372.

Baumeister RF, Stillwell AM, Heatherton TF. 1994. Guilt: An interpersonal approach. Psychol Bull. 115:243.

Bicchieri C. 2005. The grammar of society: The nature and dynamics of social norms. Cambridge University Press.

Bijsterbosch JD, Ansari TL, Smith S, Gauld O, Zika O, Boessenkool S, Browning M, Reinecke A, Bishop SJ. 2018. Stratification of MDD and GAD patients by resting state brain connectivity predicts cognitive bias. NeuroImage Clin. 19:425–433.

Blair RJR. 2013. The neurobiology of psychopathic traits in youths. Nat Rev Neurosci. 14:786.

Boonin, L. (1983). Guilt, shame and morality. The Journal of Value Inquiry, 17(4), 295–304.

Chang LJ, Gianaros PJ, Manuck SB, Krishnan A, Wager TD. 2015. A sensitive and specific neural signature for picture-induced negative affect. PLoS Biol. 13:e1002180.

Chang LJ, Smith A. 2015. Social emotions and psychological games. Curr Opin Behav Sci. 5:133–140.

Cui F, Abdelgabar AR, Keysers C, Gazzola V. 2015. Responsibility modulates pain-matrix activation elicited by the expressions of others in pain. Neuroimage. 114:371–378.

Druzgal TJ, D’Esposito M. 2001. Activity in fusiform face area modulated as a function of working memory load. Cogn Brain Res. 10:355–364.

Eisenbarth H, Chang LJ, Wager TD. 2016. Multivariate brain prediction of heart rate and skin conductance responses to social threat. J Neurosci. 36:11987–11998.

Ellsworth PC, Smith CA. 1988. From appraisal to emotion: Differences among unpleasant feelings. Motiv Emot. 12:271–302.

Fiske, A. P. 2019. The Lexical Fallacy in Emotion Research: Mistaking Vernacular Words for Psychological Entities. Psychological Review, online first publication, http://dx.doi.org/10.1037/rev0000174

Fourie MM, Thomas KGF, Amodio DM, Warton CMR, Meintjes EM. 2014. Neural correlates of experienced moral emotion: An fMRI investigation of emotion in response to prejudice feedback. Soc Neurosci. 9:203–218.

Friedman J, Hastie T, Tibshirani R. 2001. The elements of statistical learning. Springer series in statistics New York.

Frijda NH. 1993. The place of appraisal in emotion. Cogn Emot. 7:357–387.

Hamann S. 2012. Mapping discrete and dimensional emotions onto the brain: controversies and consensus. Trends Cogn Sci. 16:458–466.

Hoffman ML. 2001. Empathy and moral development: Implications for caring and justice. Cambridge University Press.

Hurtado de Mendoza A, Fernández-Dols JM, Parrott WG, Carrera P. 2010. Emotion terms, category structure, and the problem of translation: The case of shame and vergüenza. Cogn Emot. 24:661–680.

Huys QJM, Maia T V, Frank MJ. 2016. Computational psychiatry as a bridge from neuroscience to clinical applications. Nat Neurosci. 19:404.

Kédia G, Berthoz S, Wessa M, Hilton D, Martinot J-L. 2008. An Agent Harms a Victim: A Functional Magnetic Resonance Imaging Study on Specific Moral Emotions. J Cogn Neurosci. 20:1788–1798.

Koban L, Corradi-Dell’Acqua C, Vuilleumier P. 2013. Integration of error agency and representation of others’ pain in the anterior insula. J Cogn Neurosci. 25:258–272.

Koban L, Jepma M, López-Solà M, Wager TD. 2019. Different brain networks mediate the effects of social and conditioned expectations on pain. Nat Commun. 10:1–13.

Koban L, Pourtois G. 2014. Brain systems underlying the affective and social monitoring of actions: an integrative review. Neurosci Biobehav Rev. 46:71–84.

Kober H, Barrett LF, Joseph J, Bliss-Moreau E, Lindquist K, Wager TD. 2008. Functional grouping and cortical–subcortical interactions in emotion: a meta-analysis of neuroimaging studies. Neuroimage. 42:998–1031.

Kragel PA, Knodt AR, Hariri AR, LaBar KS. 2016. Decoding spontaneous emotional states in the human brain. PLoS Biol. 14:e2000106.

Kragel PA, Koban L, Barrett LF, Wager TD. 2018. Representation, pattern information, and brain signatures: from neurons to neuroimaging. Neuron. 99:257–273.

Kragel PA, LaBar KS. 2015. Multivariate neural biomarkers of emotional states are categorically distinct. Soc Cogn Affect Neurosci. 10:1437–1448.

Krishnan A, Woo C-W, Chang LJ, Ruzic L, Gu X, Lopez-Sola M, Jackson PL, Pujol J, Fan J, Wager TD. 2016. Somatic and vicarious pain are represented by dissociable multivariate brain patterns. Elife. 5:e15166.

Lepron E, Causse M, Farrer C. 2015. Responsibility and the sense of agency enhance empathy for pain. Proc R Soc B Biol Sci. 282:20142288.

Lewis-Peacock JA, Postle BR. 2008. Temporary activation of long-term memory supports working memory. J Neurosci. 28:8765–8771.

Lindquist KA, Barrett LF. 2012. A functional architecture of the human brain: emerging insights from the science of emotion. Trends Cogn Sci. 16:533–540.

Lindquist KA, Wager TD, Kober H, Bliss-Moreau E, Barrett LF. 2012. The brain basis of emotion: a meta-analytic review. Behav Brain Sci. 35:121.

Moors A, Ellsworth PC, Scherer KR, Frijda NH. 2013. Appraisal theories of emotion: State of the art and future development. Emot Rev. 5:119–124.

Pessoa L. 2017. A network model of the emotional brain. Trends Cogn Sci. 21:357–371.

Polyn SM, Natu VS, Cohen JD, Norman KA. 2005. Category-specific cortical activity precedes retrieval during memory search. Science (80-). 310:1963–1966.

Ratcliffe M. 2014. Experiences of depression: A study in phenomenology. OUP Oxford.

Redcay, E., & Schilbach, L. (2019). Using second-person neuroscience to elucidate the mechanisms of social interaction. Nature Reviews Neuroscience, 1.

Saarimäki H, Ejtehadian LF, Glerean E, Jääskeläinen IP, Vuilleumier P, Sams M, Nummenmaa L. 2018. Distributed affective space represents multiple emotion categories across the human brain. Soc Cogn Affect Neurosci. 13:471–482.

Scherer KR. 2009. Emotions are emergent processes: they require a dynamic computational architecture. Philos Trans R Soc B Biol Sci. 364:3459–3474.

Skerry AE, Saxe R. 2015. Neural representations of emotion are organized around abstract event features. Curr Biol. 25:1945–1954.

Stearns DC, Parrott WG. 2012. When feeling bad makes you look good: Guilt, shame, and person perception. Cogn Emot. 26:407–430.

Sznycer D. 2018. Forms and Functions of the Self-Conscious Emotions. Trends Cogn Sci.

Tangney JP, Stuewig J, Mashek DJ. 2007. Moral emotions and moral behavior. Annu Rev Psychol. 58:345–372.

Tilghman-Osborne C, Cole DA, Felton JW. 2012. Inappropriate and excessive guilt: Instrument validation and developmental differences in relation to depression. J Abnorm Child Psychol. 40:607–620.

Tooby J, Cosmides L. 2008. The evolutionary psychology of the emotions and their relationship to internal regulatory variables.

Tracy JL, Robins RW. 2006. Appraisal Antecedents of Shame and Guilt: Support for a Theoretical Model. Personal Soc Psychol Bull. 32:1339–1351.

Viding E, Simmonds E, Petrides K V, Frederickson N. 2009. The contribution of callous-unemotional traits and conduct problems to bullying in early adolescence. J Child Psychol Psychiatry. 50:471–481.

Wager TD, Atlas LY, Lindquist MA, Roy M, Woo C-W, Kross E. 2013. An fMRI-based neurologic signature of physical pain. N Engl J Med. 368:1388–1397.

Wager TD, Kang J, Johnson TD, Nichols TE, Satpute AB, Barrett LF. 2015. A Bayesian model of category-specific emotional brain responses. PLoS Comput Biol. 11:e1004066.

Wagner U, N’Diaye K, Ethofer T, Vuilleumier P. 2011. Guilt-Specific Processing in the Prefrontal Cortex. Cereb Cortex. 21:2461–2470.

Woo C-W, Koban L, Kross E, Lindquist MA, Banich MT, Ruzic L, Andrews-Hanna JR, Wager TD. 2014. Separate neural representations for physical pain and social rejection. Nat Commun. 5:5380.

Woo C-W, Wager TD. 2015. Neuroimaging-based biomarker discovery and validation. Pain. 156:1379.

Woo C-W, Chang LJ, Lindquist MA, Wager TD. 2017. Building better biomarkers: brain models in translational neuroimaging. Nat Neurosci. 20:365.

Yarkoni T, Poldrack RA, Nichols TE, Van Essen DC, Wager TD. 2011. Large-scale automated synthesis of human functional neuroimaging data. Nat Methods. 8:665.

Yeung N, Nystrom LE, Aronson JA, Cohen JD. 2006. Between-task competition and cognitive control in task switching. J Neurosci. 26:1429–1438.

Yu H, Duan Y, Zhou X. 2017. Guilt in the eyes: Eye movement and physiological evidence for guilt-induced social avoidance. J Exp Soc Psychol. 71.

Yu H, Hu J, Hu L, Zhou X. 2014. The voice of conscience: neural bases of interpersonal guilt and compensation. Soc Cogn Affect Neurosci. 9:1150–1158.

Yu, H., Siegel, J. Z., & Crockett, M. J. (2019). Modeling Morality in 3 - D: Decision - Making, Judgment, and Inference. Topics in cognitive science, 11(2), 409–432.

